# Loss of the Ferripyochelin Receptor FptA Drives Reduced Cefiderocol Susceptibility and Impairs Fitness in *Pseudomonas aeruginosa* PA14

**DOI:** 10.1101/2025.09.16.676464

**Authors:** Donghoon Kang, Rodrigo P. Baptista, Cesar A. Arias, William R. Miller

## Abstract

*Pseudomonas aeruginosa* is an opportunistic human pathogen and a frequent cause of multidrug-resistant infections. This organism continues to evade antimicrobial therapy despite the clinical introduction of new anti-pseudomonal antibiotics over the past several years. One of these agents is cefiderocol (FDC), a novel siderophore-cephalosporin conjugate antibiotic that was designed to overcome both intrinsic and acquired β-lactam resistance mechanisms in *P. aeruginosa*. However, studies have demonstrated that inactivation of TonB-dependent receptors, most notably the catechol siderophore receptor *piuA* can substantially curtail the drug’s ability to permeate the bacterial outer membrane, leading to rapid development of resistance. In this study, we examined the FDC resistance mechanisms of the laboratory strain PA14. We demonstrated that inactivation of the ferripyochelin receptor FptA was a first-step mutation towards FDC resistance. Through transposon mutagenesis, we identified several resistance pathways following *fptA* inactivation, such as the loss of an additional FDC import porin and overexpression of the MuxABC-OpmB multidrug efflux system. Introduction of clinically-identified mutations analogous to these transposon insertions in the absence of *fptA* conferred full FDC non-susceptibility while preserving the activity of other antipseudomonal β-lactam antibiotics. We also demonstrated that inactivation of *fptA* in a pyoverdine biosynthetic mutant disrupted bacterial iron homeostasis and conferred a fitness disadvantage. These FDC resistance mechanisms identified in PA14 highlight the long-term challenges of using FDC treatment for drug-resistant *P. aeruginosa* infections.

## INTRODUCTION

*Pseudomonas aeruginosa* is a leading cause of healthcare associated infections, particularly in critically ill individuals and those with chronic lung diseases such as cystic fibrosis (1). Therapy for infections due to these organisms is complicated by clinical isolates displaying difficult-to-treat resistance (DTR) that includes resistance to fluoroquinolones, piperacillin-tazobactam, cefepime, ceftazidime, aztreonam, and carbapenems (2). Rates of resistance to the newer β-lactam/β-lactamase inhibitor (BL/BLI) combinations such as ceftolozane-tazobactam, ceftazidime-avibactam, and imipenem-relebactam have also increased among global collections of extensively-drug resistant isolates (3, 4). Cefiderocol (FDC) is a siderophore conjugated cephalosporin that retains *in vitro* activity against drug-resistant *P. aeruginosa* by utilizing TonB-dependent receptor iron transporters to facilitate drug uptake (5). Despite high rates of susceptibility *in vitro*, clinical failure and the emergence of resistance to FDC on therapy are growing concerns (6-8).

A number of mutations have been linked to FDC resistance from both clinical isolates before and after antibiotic exposure, as well as *in vitro* adaptation assays. These can be grouped into several categories. First, alterations of outer membrane TonB-dependent siderophore transporters or their regulatory components are postulated to act through inactivation or down-regulation of FDC sites of entry (9, 10). These changes may arise during FDC exposure, although decreased expression of the major catechol transporters *piuA* and *pirA* due to frameshift mutations in the transcriptional regulator *pirR* have been noted in the *P. aeruginosa* population prior to the introduction of FDC (11). Second, mutations leading to increased activation of the CpxS histidine kinase have been demonstrated to increase expression of two efflux systems, MexAB-OprM and MuxABC-OpmB (12, 13), possibly resulting in antibiotic efflux or increased secretion of pyoverdine (14). Finally, the presence of certain exogenous β-lactamases or mutations in the intrinsic pseudomonal AmpC cephalosporinase have been associated with FDC resistance (15, 16), and may be selected by prior exposure to ceftolozane-tazobactam or ceftazidime-avibactam (17, 18).

*P. aeruginosa* possesses two endogenous siderophores with different affinities for ferric iron, pyoverdine and pyochelin. Pyoverdine has a distinctly high affinity for ferric iron (K_d_ = 10^-32^ M) (19) and *in vitro* data suggest it may chelate the metal from FDC (5), preventing the antibiotic from utilizing TonB-dependent receptors for cell entry (14). Conversely, pyochelin has a lower iron affinity, and production of this siderophore leads to upregulation of the cognate TonB-receptor FptA (20, 21). Overexpression of FptA results in increased susceptibility to FDC, likely through increased antibiotic uptake (9), and we have previously identified *fptA* inactivating mutations in laboratory and clinical isolates that developed FDC resistance during therapy (6, 11). However, their specific contribution to the resistance phenotype has not been addressed. The aim of this study was to characterize the acquisition and impact of *fptA* mutations on the FDC susceptibility and fitness of the laboratory strain *P. aeruginosa* PA14.

## RESULTS

### fptA inactivation is a first-step mutation towards FDC resistance in PA14

Previous studies of *in vitro* adapted and clinical isolates of *P. aeruginosa* identified mutations in *fptA* associated with decreased FDC susceptibility (6, 11). To confirm whether the loss of the ferripyochelin receptor FptA contributes to cefiderocol (FDC) resistance in the laboratory reference strain PA14, we generated a *fptA* gene deletion mutant. Loss of *fptA* resulted in decreased susceptibility to FDC, indicated by a significant decrease in the antibiotic disk diffusion diameter and 2-4-fold increase in the minimum inhibitory concentration (MIC) (**Fig. 1A, B**). It was noted that isolated inner-zone colonies emerged within 48 hours on disk diffusion assay with wild type PA14 (**Fig, 1C**). These colonies were purified, passaged in drug-free medium, and evaluated for FDC susceptibility by disk diffusion testing. Nearly all (n=10/12) inner-zone colonies exhibited decreased susceptibility to FDC (**Fig. 1B, C**), indicating that these 10 colonies represented spontaneous, drug-adapted mutants as opposed to persister cells.

**Figure 1.**
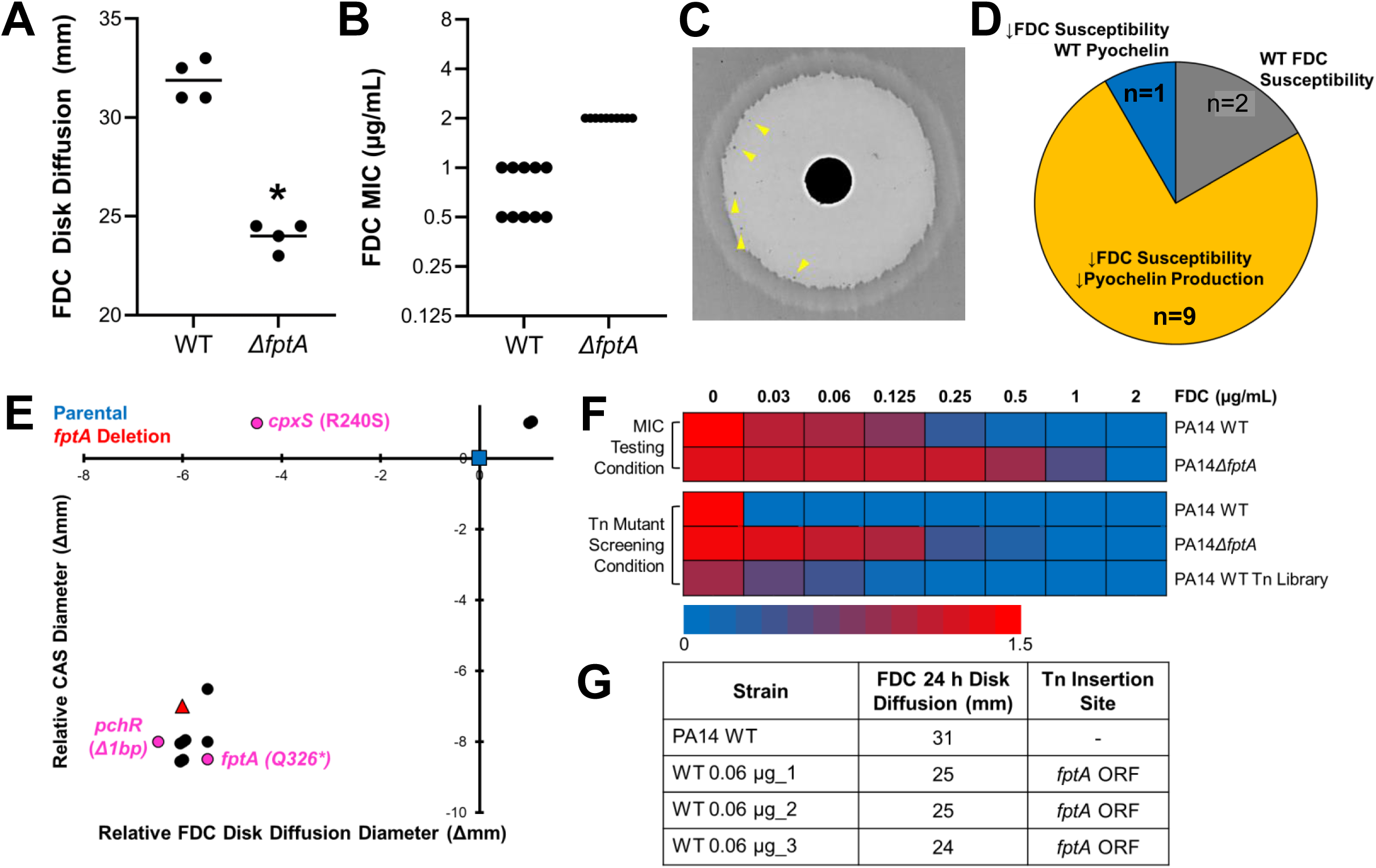
FptA loss is a first-step mutation towards FDC resistance in PA14. **(A,B)** FDC susceptibility for PA14 WT and PA14*ΔfptA*. FDC disk diffusion diameters **(A)** and MICs **(B)** were measured at 24 h. **(C)** FDC disk diffusion assay image for PA14 WT after 48 h growth. Yellow arrows point to inner zone colonies. **(D)** Summary of FDC susceptibility and pyochelin production profiles of FDC inner zone colonies from PA14 WT. **(E)** Changes in FDC susceptibility (disk diffusion diameter) and pyochelin production (Chrome Azurol S assay diameter) for inner zone colonies passaged in drug-free medium, compared to PA14 WT. Blue square: PA14 WT. Red triangle: *PA14ΔfptA*. Circles: strains from inner zone colonies. Magenta circles: strains analyzed by whole genome sequencing and their relevant mutations (full list of mutations provided in **Table S1**). **(F)** PA14 WT, PA14*ΔfptA*, or library of ∼50K transposon mutants from PA14 WT grown in increasing concentrations of FDC in iron-depleted Mueller-Hinton broth (MIC testing medium) or modified low-iron casamino acids medium (Tn mutants screening condition). Bacterial growth (O.D. 600 nm) was measured after 24 h. **(G)** FDC susceptibility (disk diffusion diameter) and transposon insertion location of three random mutants isolated from the FDC 0.06 µg/mL growth condition. * corresponds to *p* < 0.01 based on a Student’s *t*-test. Each point in **(B)** corresponds to MIC interpreted from a technical replicate from at least three biological replicates.

We next screened these spontaneous mutants for siderophore production via a Chrome Azurol S (CAS) activity assay. Removal of iron from CAS by *P. aeruginosa* siderophores causes a chromic shift (blue to yellow), producing a halo around the bacterial lawn (**Fig. S1A**) (22). On Mueller-Hinton agar, siderophore activity was primarily driven by the smaller and more diffusible siderophore pyochelin, rather than pyoverdine. The pyochelin biosynthetic mutant PA14*pchE* produced a significantly smaller apo-CAS halo compared to wild-type PA14 (**Fig. S1**). In contrast, the pyoverdine biosynthetic mutant PA14*pvdF* produced a significant larger halo, probably due to increased pyochelin production in the absence of pyoverdine. Consistent with FptA’s established role in the regulation of pyochelin production by a positive feedback loop (i.e.: import of ferripyochelin by FptA promotes siderophore biosynthesis) (20), PA14*ΔfptA* exhibited low siderophore activity on the CAS assay.

We took advantage of the results of the CAS assay above to screen inner-zone colonies for *fptA* loss-of-function mutations. A total of 9/10 FDC-adapted spontaneous mutants derived for the FDC inner zone exhibited poor pyochelin production (**Fig. 1D, E**). To confirm the inactivation of *fptA* expression, we performed whole-genome sequencing and mutation analysis. Of the two mutants with decreased pyochelin production sequenced, one harbored a nonsense mutation in *fptA* while the other harbored a frameshift mutation in *pchR*, which encodes the transcriptional regulator for *fptA*. None of these mutants harbored additional mutations previously associated with cefiderocol resistance (**Table S1**). The one mutant with decreased FDC susceptibility, but wild-type pyochelin production, was found to have a mutation in *cpxS*. This preference for *fptA* inactivation or downregulation in FDC-adapted mutants indicated that FptA loss was likely a first-step towards FDC resistance in PA14.

To investigate if *fptA* was also selectively targeted during a single FDC exposure in iron-depleted media, we performed a transposon insertion screen in *P. aeruginosa* PA14, generating a library of ∼50K insertion mutants with the MAR2xT7 mariner transposon (23). Three random mutants were isolated from the batch transposon library culture with the highest concentration of FDC that permitted bacterial growth (**Fig. 1F**). Transposon insertion sites were identified by Sanger sequencing. All mutants exhibited decreased FDC susceptibility by disk diffusion testing and all harbored transposon insertions in the *fptA* open reading frame (**Fig. 1G**), supporting the conclusion that FptA loss was a first-step adaptation to FDC exposure.

### Second step mutations following fptA-inactivation lead to FDC nonsusceptibility in PA14

Previously, we have identified second step mutations as an important driver of FDC non-susceptibility in clinical isolates (11). To systematically identify additional mutations associated with loss of FDC susceptibility in the absence of FptA, we performed transposon mutagenesis in PA14*ΔfptA* generating ∼50,000 insertion mutants that were pooled into 12 sub-libraries. Each sub-library was grown in iron-depleted medium with increasing concentrations of FDC, and at least three random mutants were isolated from the culture with the highest concentration of FDC that permitted bacterial growth (**Fig. 2A**). We identified three distinct mutants with at least a two-fold increase in the FDC MIC compared to the *ΔfptA* parental strain (**Fig. 2B**). These insertions were in open reading frames of *piuA* (encoding a catechol siderophore receptor), *PA14_31850* (encoding a hypothetical protein), and *pilM* (type IV pilus biosynthesis gene). While *piuA* encodes a previously identified TonB receptor important for FDC import (9), the latter two insertion sites were not previously characterized in FDC or β-lactam resistance genes. Each insertion site was immediately upstream of *muxA* (multidrug efflux pump gene) or *ponA* (encoding penicillin-binding-protein 1A, PBP1A), respectively. The MAR2xT7 mariner transposon was previously designed to not affect the expression of potential downstream essential genes by allowing transcription from the aminoglycoside resistance cassette within the transposon (23). We thus hypothesized that the transposon insertions upregulated expression of *muxA* and *ponA*. We measured the mRNA levels of these downstream genes by qRT-PCR, which confirmed their overexpression in the transposon mutants (**Fig. 2C**).

**Figure 2.**
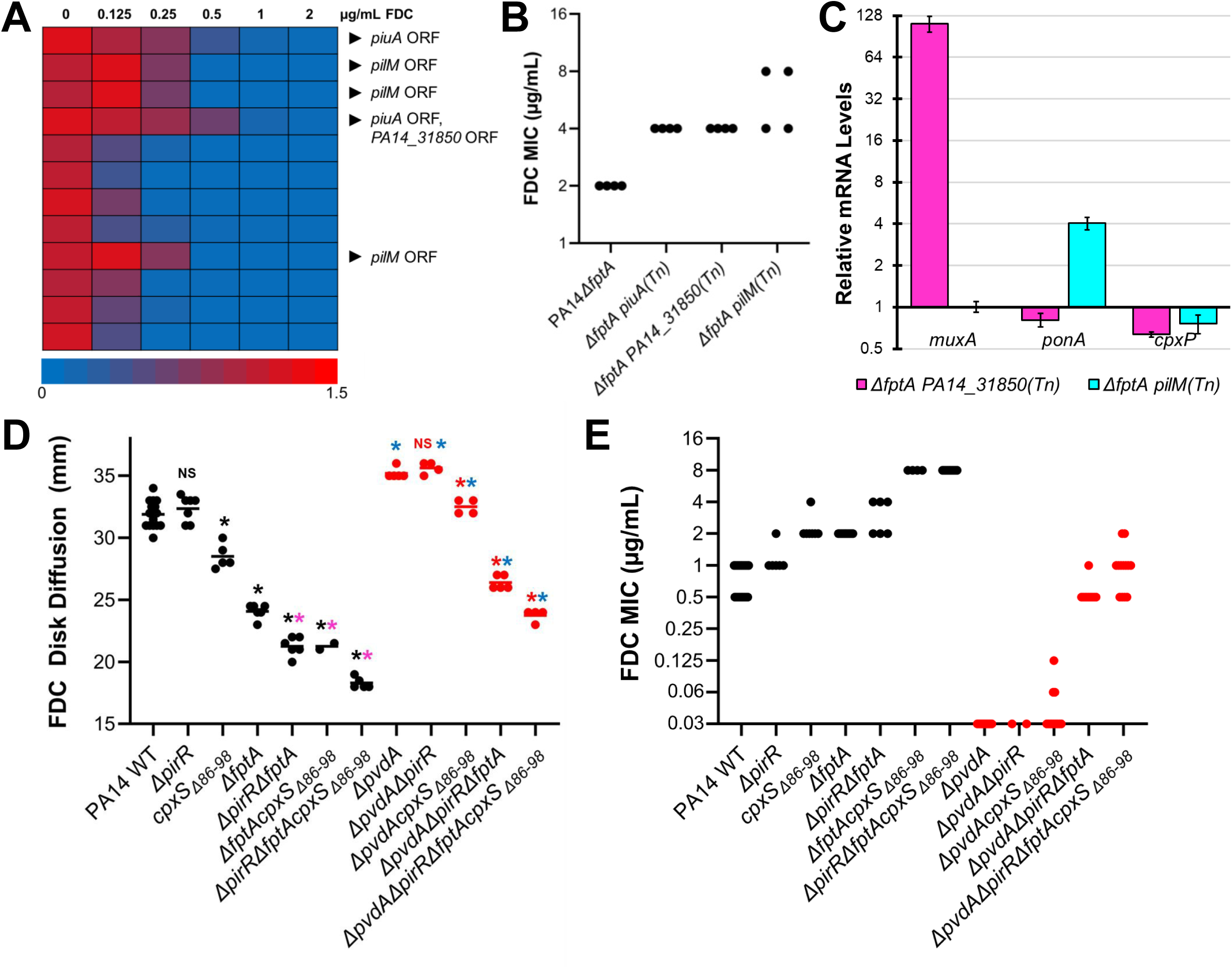
Clinically identified mutations confer full FDC nonsusceptibility in PA14. **(A)** Library of ∼50K transposon mutants from PA14*ΔfptA*, split into twelve sublibraries, grown in increasing concentrations of FDC in iron-depleted broth medium. Bacterial growth (O.D. 600 nm) was measured after 24 h. Black arrows point to transposon insertion locations from mutants that were isolated from the condition with the highest concentration of FDC that permitted bacterial growth. **(B)** FDC MICs for PA14*ΔfptA* and three transposon mutants, *piuA (Tn)*, *PA14_31850 (Tn)*, and *pilM (Tn)*, determined after 24 h growth. **(C)** Relative expression of *muxA*, *ponA*, and *cpxP* (by qRT-PCR) in PA14*ΔfptA* mutants with transposon insertions in *PA14_31850* or *pilM* compared to the PA14*ΔfptA* parental strain. **(D, E)** FDC susceptibility for PA14 mutants. FDC disk diffusion diameters **(D)** and MICs **(E)** were measured at 24 h.* corresponds to *p* < 0.01, NS corresponds to *p* > 0.05 based on a one-way ANOVA with Tukey’s multiple comparisons test. Black: compared to PA14 WT. Magenta: compared to PA14*ΔfptA*. Red: compared to PA14*ΔpvdA*. Blue: comparing *ΔpvdA* mutants to their pyoverdine-producing counterparts. Each point in **(B, E)** corresponds to MIC interpreted from a technical replicate from at least two biological replicates.

We have previously identified combinations of *pirR* inactivation, *fptA* inactivation, and increased *cpxS* activity in FDC-nonsusceptible clinical isolates, although their specific contribution to the phenotype was not clear (6, 11). Based on these findings and the transposon insertion data, we sought to confirm that introduction of these clinically identified mutations was responsible for the changes in susceptibility to FDC.

Single deletion mutants were constructed for PA14*ΔpirR* in addition to replacing the *cpxS* allele with a variant resulting in deletion of amino acids 86 to 98 as identified in a resistant clinical isolate (PA14*cpxS*_Δ86-98_, **Fig. S2A**) (6). This mutation occurs in the periplasmic sensor domain of CpxS and leads to increased expression of *muxA* and another CpxR-target gene, *cpxP* (**Fig. S2B**). The FDC disk diffusion diameter and MIC for the PA14*ΔpirR* strain was not significantly different from PA14 wild type (**Fig. 2D, E**). The PA14*cpxS*_Δ86-98_ strain showed a decrease in FDC disk diffusion diameter and increase in MIC of ∼2-fold, consistent with altered FDC susceptibility. Multi-gene variants demonstrated further decreases in FDC susceptibility, with the FDC MIC for the PA14*ΔfptAΔpirR* double mutant increasing to near the CLSI breakpoint of 4 μg/mL and the PA14*ΔfptAcpxS*_Δ86-98_ displaying a non-susceptible MIC of 8 μg/mL.

Interestingly, PA14*cpxS_Δ86-98_* and PA14*ΔfptA* had similar FDC MICs (2 µg/mL) despite our observations that FptA loss was predominantly favored as a first-step adaptation to FDC exposure. To compare the impact of these mutations, we measured the bacterial growth kinetics in increasing concentrations of FDC (**Fig. S3**). The WT strain exhibited severely delayed growth at FDC concentrations as low as 0.03 µg/mL. Mutations in *pirR* or *cpxS* showed similar growth profiles with an increase in lag phase. In contrast, deletion of *fptA* resulted in less pronounced growth deficits at low FDC concentrations, suggesting improved fitness in the presence of FDC.

While mutations in *cpxS* have been broadly associated with β-lactam resistance (14), mutations in TonB-dependent siderophore receptors or their regulatory elements, such as *ΔfptA* or *ΔpirR*, would be predicted to only affect the import of FDC. To test this hypothesis, we performed antibiotic susceptibility testing for various β-lactams and β-lactam/β-lactamase-inhibitor combinations that are used to treat multidrug-resistant *P. aeruginosa*. Mutations exclusive to siderophore import did not alter the MIC of other β-lactams or combination agents, with no change between PA14 and PA14*ΔfptAΔpirR* double mutant for imipenem-relebactam, ceftazidime-avibactam, ceftolozane-tazobactam, cefepime, or aztreonam (**Table 1**). Compared to WT PA14, mutants with the *cpxS_Δ86-98_*allele exhibited a modest ∼2-fold increase in the MICs for ceftazidime-avibactam, ceftolozane-tazobactam, and aztreonam, although these MICs remained within the susceptible range. These results demonstrate that while clinically-identified FDC resistance mechanisms were sufficient to confer FDC nonsusceptibility in the laboratory strain PA14, the activity of other agents reserved for drug-resistant *P. aeruginosa* infections was minimally impacted.

**Table 1.** Cefiderocol Resistance Mechanisms Preserve the Activity of Other Antipseudomonal β-Lactam Antibiotics. MICs for antipseudomonal β-lactams and β-lactam/β-lactamase inhibitor combinations for PA14 WT, PA14*ΔpirRΔfptA*, PA14*cpxS_Δ86-98_*, and PA14*ΔpirRΔfptAcpxS_Δ86-98_*. MICs were determined by gradient diffusion strip test on Mueller-Hinton agar after 18 h.

| MICs (µg/mL) | Imipenem-relebactam |  | Ceftazidime-avibactam |  | Ceftolozane-tazobactam |  | Cefepime |  | Aztreonam |  |
| --- | --- | --- | --- | --- | --- | --- | --- | --- | --- | --- |
|  | Range | Mode | Range | Mode | Range | Mode | Range | Mode | Range | Mode |
| PA14 WT | 0.38-0.5 | 0.5 | 0.5-1 | 0.5 | 0.38-0.5 | 0.5 | 1 | 1 | 2-3 | 3 |
| PA14Δ*pirR*Δ*fptA* | 0.38-0.5 | 0.5 | 0.5-0.75 | 0.5 | 0.38 | 0.38 | 1-1.5 | 1 | 3 | 3 |
| PA14*cpxS*Δ<sub>86-98</sub> | 0.38-0.5 | 0.38 | 1.5 | 1.5 | 1-1.5 | 1.5 | 1-1.5 | 1.5 | 6 | 6 |
| PA14Δ*pirR*Δ*fptA**cpxS*Δ<sub>86-98</sub> | 0.38-0.5 | 0.38 | 1-1.5 | 1 | 1-2 | 1.5 | 1 | 1 | 4-6 | 6 |

### Impact of pyoverdine production on FDC susceptibility

In addition to decreased import via loss of siderophore uptake receptors, pyoverdine production has been implicated in FDC resistance in *P. aeruginosa* (14). Several reports have demonstrated a positive correlation between pyoverdine production and FDC resistance in *P. aeruginosa* clinical isolates (14, 24). To determine the extent of pyoverdine’s role in the FDC-nonsusceptible mutant PA14Δ*pirRΔfptAcpxS_Δ86-98_*, we introduced these mutations into PA14*ΔpvdA*, a pyoverdine biosynthetic mutant. In the absence of pyoverdine, the three FDC resistance mechanisms were not sufficient for full nonsusceptibility (MIC=1 µg/mL) (**Fig. 2E**). We observed similar ∼8-fold decreases in FDC MICs and significant increases in FDC disk diffusion diameters for each intermediate mutant we generated in the *ΔpvdA* background (**Fig. 2D, E**), supporting the role of pyoverdine production in FDC resistance.

### Loss of FptA confers a fitness cost in a pyoverdine biosynthetic mutant

We posited that the inability to uptake ferripyochelin in the absence of pyoverdine production would substantially disrupt bacterial iron homeostasis, conferring a fitness cost in iron-restricted media. To test this hypothesis, we measured bacterial growth kinetics in IDMH with or without ferric iron supplementation (100 µM FeCl_3_) for pyoverdine-producing (WT PA14, PA14*ΔpirR*, PA14*ΔpirRΔfptA*) and non-producing (PA14*ΔpvdA*, PA14*ΔpvdAΔpirR*, PA14*ΔpvdAΔpirRΔfptA*) mutants. In pyoverdine-producing strains, inactivation of *fptA* did not affect bacterial growth in IDMH (**Fig. 3A**). In the *ΔpvdA* mutants, loss of *fptA* hampered bacterial growth (**Fig. 3B**), with significantly lower densities (O.D. 600 nm) throughout the growth curve (**Table S2**). However, these growth defects were fully rescued with ferric iron supplementation (**Fig. 3C, Table S2**), demonstrating that iron starvation impeded bacterial growth in the absence of pyoverdine production and ferripyochelin uptake.

**Figure 3.**
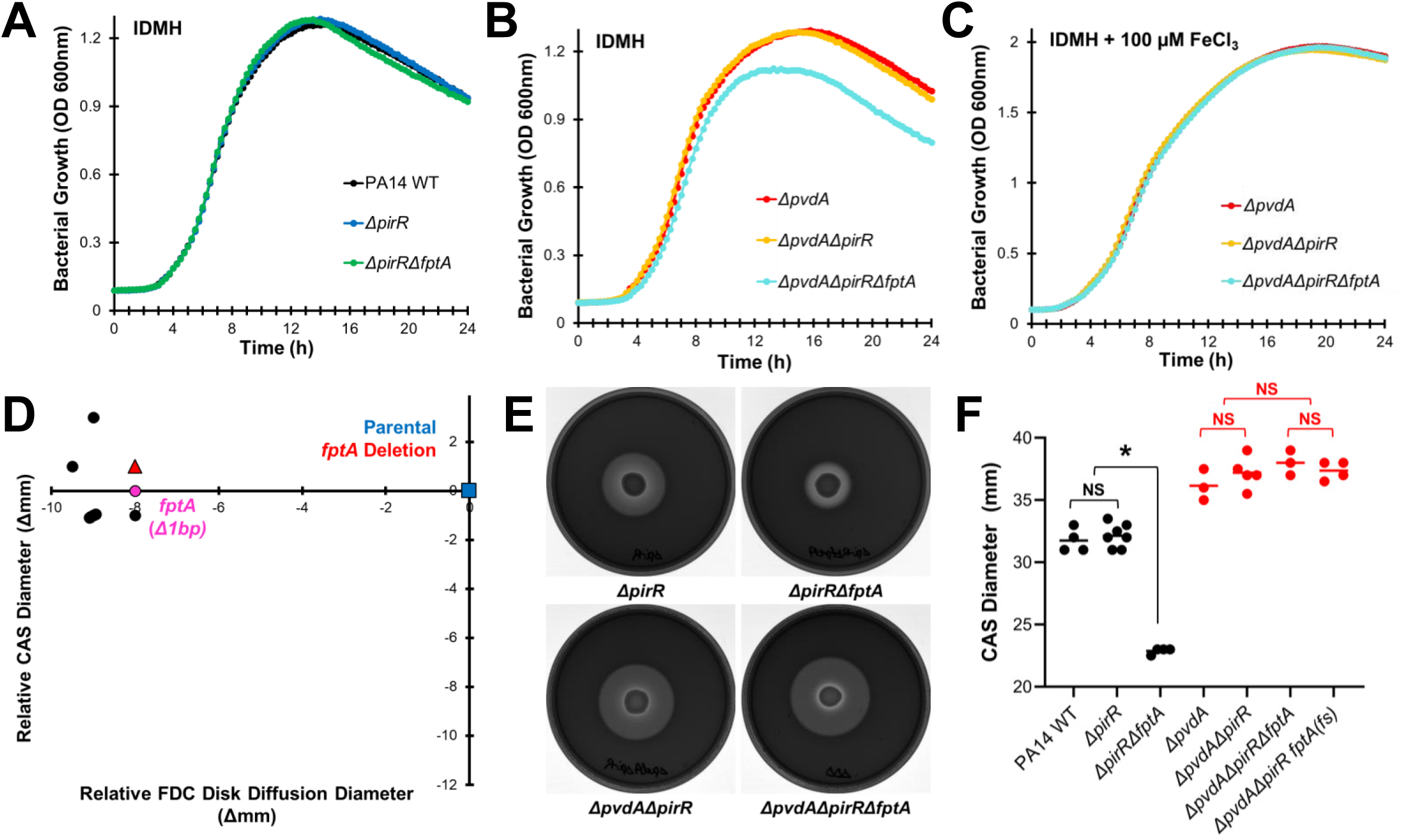
Inactivation of *fptA* confers a fitness cost in a pyoverdine biosynthetic mutant. **(A-C)** Bacterial growth (O.D. 600 nm) measured every 30 min for 24 h in iron-depleted Mueller-Hinton broth with **(A, B)** or without **(C)** ferric iron (100 µM FeCl_3_) supplementation for PA14 mutants. Each point represents the average across 4 biological replicates. **(D)** Changes in FDC susceptibility (disk diffusion diameter) and pyochelin production (Chrome Azurol S assay diameter) for inner zone colonies isolated from a FDC disk diffusion assay for PA14*ΔpvdAΔpirR*, compared to parental strain control. Blue square: PA14*ΔpvdAΔpirR*. Red triangle: PA14*ΔpvdAΔpirRΔfptA*. Circles: strains from inner zone colonies. Magenta circle: strain analyzed by whole genome sequencing, relevant mutation indicated (full list of mutations provided in **Table S1**). **(E)** Visualization of pyochelin production by PA14 mutants on Mueller-Hinton agar supplemented with CAS-Fe^3+^ after 48 h. **(F)** Diameter of the apo-CAS halo that represents pyochelin production. * corresponds to *p* < 0.01, NS corresponds to *p* > 0.05 based on a one-way ANOVA with Tukey’s multiple comparisons test.

Based on these findings, we examined whether FptA loss remained favored as an adaptation strategy towards FDC resistance in the pyoverdine biosynthetic mutant PA14*ΔpvdAΔpirR*. We screened inner-zone colonies that emerged within 48 h from a FDC disk diffusion assay, verifying decreased FDC susceptibility for the spontaneous mutants by disk diffusion testing and reduced pyochelin production using a CAS siderophore activity assay (**Fig. 3D**). None of the FDC-adapted mutants exhibited decreased pyochelin production. We performed whole-genome sequencing and mutation analysis for one randomly selected mutant. Unexpectedly, this mutant harbored a *fptA* frameshift mutation with no other mutations previously associated with FDC resistance (**Table S1**). Measuring pyochelin production in the PA14*ΔpvdAΔpirRΔfptA* deletion mutant by CAS assay confirmed that loss of the ferripyochelin receptor in the absence of pyoverdine did not downregulate pyochelin biosynthesis (**Fig. 3E, F**), possibly due to a severe iron-starvation response. Without functional FptA, this mutant would not have been able to effectively utilize pyochelin, which could have also contributed to the growth defects we observed. Importantly, while we were not able to fully screen FDC-adapted, pyoverdine biosynthetic mutants for the loss of *fptA* expression or function, the selection of the *fptA* frameshift mutant suggests that inactivation of this outer membrane porin remained one of the major mechanisms of FDC resistance despite the fitness cost.

### Fitness costs drive fptA allele frequency in mixed populations

Finally, we hypothesized that in the absence of pyoverdine, acquisition of *fptA* mutations could result in unstable subpopulations with reduced FDC susceptibility. Without selective pressure from FDC to offset fitness costs, these mutations would not be maintained at stable frequencies across the population. As proof of concept, we passaged inoculum-controlled (identical starting inoculum of ∼10^6^ CFU/mL, **Fig. 4A**) co-cultures of isogenic *fptA*^+^ and *fptA*^-^ strains in the presence (PA14*ΔpirR*, PA14*ΔpirRΔfptA*) or absence (PA14*ΔpvdAΔpirR*, PA14*ΔpvdAΔpirRΔfptA*) of pyoverdine production. Throughout the 10-day experiment, FDC susceptibility of the pyoverdine-producing co-cultures remained stable, with no significant changes in FDC disk diffusion diameters between day 1 and 10 of passaging (**Fig. 4B-D**). However, co-cultures of the pyoverdine biosynthetic mutants exhibited a gradual sensitization of the population to FDC with a visible dwindling of the *ΔfptA* subpopulation, which was no longer observable by the end of the experiment (**Fig. 4B-D, Fig. S4**). These observations were validated by qPCR analysis, which indicated a significant, 3-log reduction in the frequency of the *ΔfptA* allele between the first and tenth passage for the pyoverdine-null cultures (**Fig. 4E**). In contrast, the *ΔfptA* allele remained stable for the pyoverdine-producing cultures. These results suggest that for *P. aeruginosa* strains that do not produce pyoverdine, the selection of *fptA* mutations may rely on continuous selective pressure from the antibiotic and represent an unstable subpopulation with reduced FDC susceptibility.

**Figure 4.**
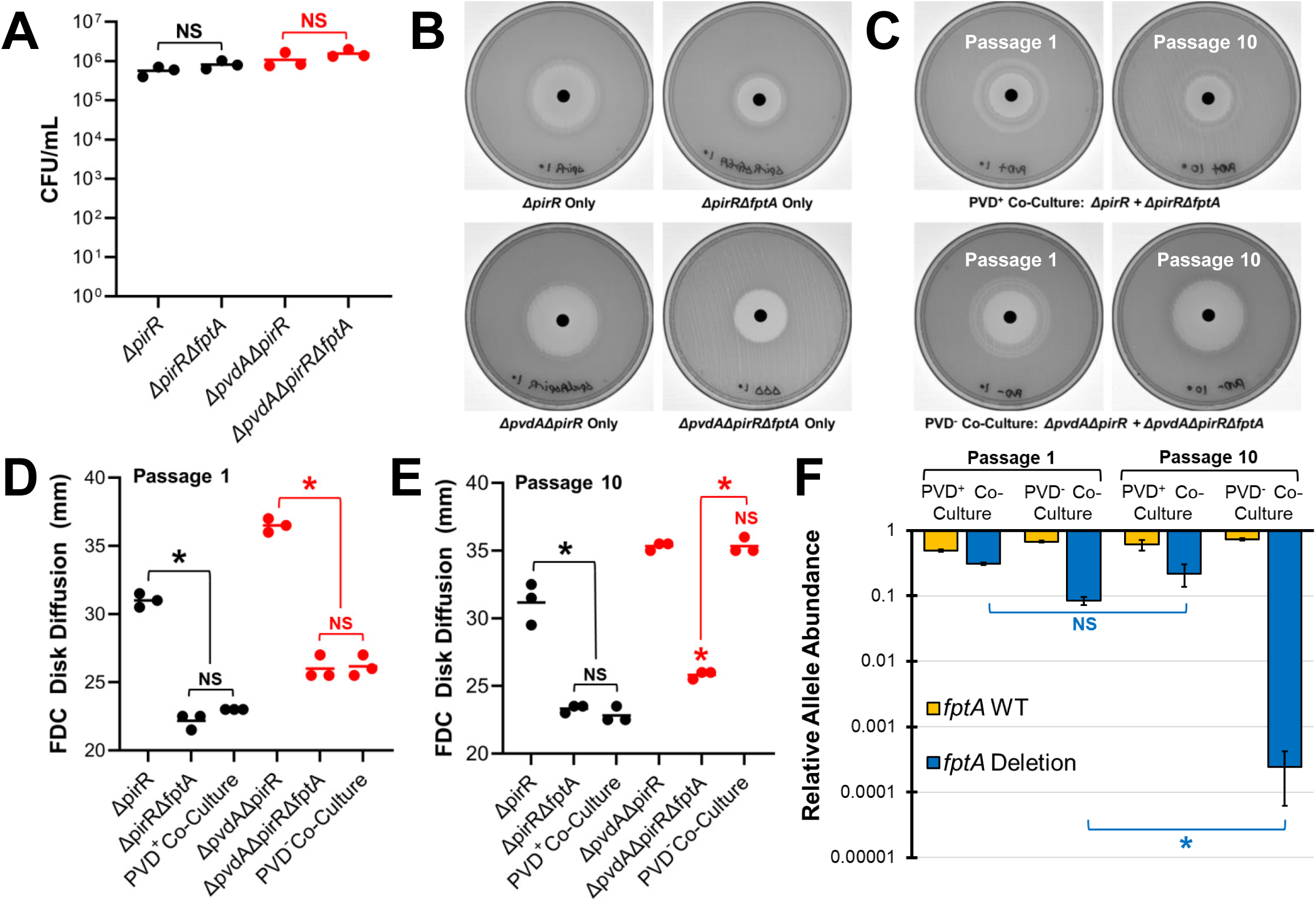
*ΔfptA* is not stably maintained in a pyoverdine biosynthetic mutant. **(A)** Initial inoculum of PA14 mutants in the serial passage experiment. **(B)** Images of FDC disk diffusion testing plates for monocultures of PA14 mutants. **(C)** Images of FDC disk diffusion testing plates for co-cultures of PA14*ΔpirR* and PA14*ΔpirRΔfptA* (top) or PA14*ΔpvdAΔpirR* and PA14*ΔpvdAΔpirRΔfptA* (bottom) passaged in iron-depleted Mueller-Hinton broth for 1 or 10 days. **(D, E)** Quantification of FDC disk diffusion diameters from assays performed after the first **(D)** or tenth passage **(E)**. **(F)** Quantification of the *fptA* WT and *ΔfptA* alleles in the co-cultures after the first or tenth passage via qPCR using allelic-specific primers. * corresponds to *p* < 0.01, NS corresponds to *p* > 0.05 based on a one-way ANOVA with Sidak’s **(A, E)** or Tukey’s **(C, D)** multiple comparisons test.

## DISCUSSION

Mechanisms of resistance to FDC are complex and result from an overlapping number of pathways that may alter siderophore import, iron homeostasis, efflux pumps, and hydrolysis via β-lactamases (25). Genomic data from *in vitro* adaptation experiments and clinical isolates collected prior to the introduction of FDC suggest there may be distinct pathways by which FDC resistance emerges depending on the selective pressure. Under direct FDC pressure, mutations in TonB receptor pathways (PirR, PiuA, FptA) or CpxS are frequently found (6, 11, 14, 15, 26, 27). Exposure to newer β-lactam/β-lactamase inhibitor combinations including ceftolozane-tazobactam and ceftazidime-avibactam have been associated with alterations in the intrinsic AmpC cephalosporinase which may predispose to FDC resistance in the setting of decreased uptake via TonB receptor loss (17, 18, 28). Thus, it is important to understand pathways to the emergence of FDC resistance to help guide selection of therapeutics for DTR-*P. aeruginosa*.

In this study, we used the laboratory strain PA14 to characterize the stepwise events leading to FDC non-susceptibility. We found that loss of function of the ferripyochelin receptor FptA was a frequent first step to reduced susceptibility on both Mueller-Hinton agar and in iron-depleted broth medium. This is contrary to a previous study that examined the laboratory strain PAO1, where the inactivation of the catechol siderophore receptor *piuA* appeared to be the first-step mutation towards FDC resistance (15). In another PAO1-based study, disruption of *fptA* by transposon insertion did not affect the FDC MIC, while disruption of *piuA* increased the MIC by 16-fold (5). Similarly, while some *P. aeruginosa* strains exhibited loss of *piuA* under *in vivo* or *in vitro* FDC selective pressure (26, 27), others exhibited loss of *fptA* (6, 11, 29), both (11, 15), or neither (27), indicating that inactivation of these TonB-dependent receptors may have disparate impacts on FDC import depending on the *P. aeruginosa* strain (30).

In PA14, single mutations of *fptA* were not sufficient to confer a resistant phenotype and we identified a number of second step mutations that work in concert with decreased uptake via loss of FptA. Several of these mechanisms have been previously described, including upregulation of genes in the *cpxS* regulon (14) and inactivation of the siderophore transporter encoded by *piuA* (5, 9, 31). However, we also identified a transposon insertion mutant that led to increased expression of the *ponA* gene encoding PBP1A. Increased levels of the bifunctional peptidoglycan transpeptidase/glycosyltransferase PBP1A may partially compensate for the inhibition of PBP3-transpeptidase activity by FDC (32).

While previous studies have identified mutations in *pirR*, *cpxS*, and *fptA* in association with FDC resistance, these mutations were frequently found in conjunction with multiple other changes making discerning their specific contribution difficult (14, 15). This study compared the direct impact of these mutations alone and in combination using allelic exchange in a laboratory *P. aeruginosa* background, confirmed via whole genome sequencing. Individually, loss of FptA and activation of CpxS both led to a shift in FDC MIC, and together these changes were sufficient to confer an FDC non-susceptible phenotype. In contrast, loss of PirR showed minimal changes in FDC disk diffusion diameter or MIC, suggesting additional differences in growth environment or strain background are needed for the loss of PirR to exert an effect. Importantly, we found that mutations arising from FDC exposure in PA14 display limited cross-resistance with other anti-pseudomonal β-lactam antibiotics, including imipenem-relebactam, ceftazidime-avibactam, and ceftolozane-tazobactam. This contrasts with the potential for the selection of cross-resistance to FDC that has been observed after ceftolozane-tazobactam or ceftazidime-avibactam exposure in the literature (17, 18, 33). While further study is needed, this lack of cross-resistance after FDC exposure could suggest the use of FDC up front against select DTR-*P. aeruginosa* isolates, rather than as a salvage regimen.

Interestingly, overexpression of the *muxABC*-*opmB* operon via *cpxS_Δ86-98_* reduced bacterial susceptibility to FDC even in the absence of pyoverdine (in PA14*ΔpvdAcpxS_Δ86-98_* and PA14*ΔpvdAΔpirRΔfptAcpxS_Δ86-98_* backgrounds). This contrasts with a previous study that hypothesized that mutations in *cpxS* contribute to FDC resistance in a pyoverdine-dependent manner, where MuxABC-OpmB is predicted to have a direct role in pyoverdine secretion (14). In iron-depleted Mueller-Hinton broth (IDMH), the *cpxS_Δ86-98_* mutation did not affect pyoverdine production, suggesting that increased drug efflux may be responsible for the decrease in FDC susceptibility observed in our study.

Finally, we determined the fitness of *fptA* deletion mutants *in vitro* across both iron-limited conditions and in strains deficient in pyoverdine production. Our results suggest that pyoverdine production mitigates the impact of FptA loss, likely by providing an alternative pathway for iron acquisition. The concomitant production of pyoverdine with loss of ferripyochelin import may synergize in maintaining clones with reduced FDC susceptibility in the population. Indeed, in a pyoverdine biosynthetic mutant background, passage in the absence of FDC led to a decrease in the frequency of the *fptA* deletion allele. The resulting population demonstrated a susceptible disk diffusion diameter despite a low frequency of *fptA* mutants in the population, suggesting a possible mechanism for the emergence of FDC heteroresistance in *P. aeruginosa* strains that have lost the ability to produce pyoverdine, commonly found in cystic fibrosis patients (34-37).

In summary, we evaluated the major contributions of FptA to FDC susceptibility and fitness in the laboratory strain PA14. Loss of FptA along with second step mutations in the CpxS histidine kinase, TonB receptors, and increased expression of PBP1A were found to contribute to decreased FDC activity. Strains with deletion of *fptA* demonstrated fitness defects in the absence of pyoverdine production, which contributed to a decrease in the allele frequencies of the deletion mutant in a co-culture experiment. Further work is needed to explore the intersection of FDC-resistance associated fitness costs and the heteroresistant phenotype.

## MATERIALS AND METHODS

### Bacterial Strains and Allelic Exchange Mutagenesis

The *ΔpirR* (Δ444 bp coding), *ΔfptA* (Δ1,163 bp coding), and *cpxS*_Δ86-98_ mutations were introduced into *P. aeruginosa* PA14 (38) and pyoverdine biosynthetic mutant PA14*ΔpvdA* (39) using allelic exchange mutagenesis by the pEXG2 vector as previously described (40). In brief, regions upstream and downstream (∼600 bp) of the gene deletion site were amplified by polymerase chain reaction (PCR) and cloned into linearized pEXG2 (digested by SacI and XbaI (New England Biolabs, NEB)) via Gibson assembly (NEB). The assembled vector was transformed into competent *Escherichia coli* DH5α cells (NEB) by selecting for gentamicin resistance. The tandem insertion of the upstream and downstream regions into the plasmid was verified by PCR and Sanger sequencing (Azenta Genewiz). The vector was then transformed into competent *E. coli* SM10 conjugal donor cells. PA14 and SM10 were mated on Tryptic Soy Agar (TSA) (BD) and single-crossover merodiploid mutants were selected on TSA supplemented with 30 µg/mL gentamicin (Millipore Sigma) and 10 µg/mL triclosan (Millipore Sigma). Merodiploid colonies were grown in antibiotic-free LB broth (BD) and counter-selected for the *sacB* gene on no sodium LB agar (NSLB) with sucrose (5 g/L yeast extract (BD), 10 g/L tryptone (Gibco), 20 g/L agar (BD), 20% w/v sucrose (Fisher Scientific), 10 µg/mL triclosan). PA14 mutants were verified by PCR to confirm the gene deletion and by Illumina sequencing to confirm the absence of additional extraneous mutations. All primer sequences are available in **Table S3**.

### Cefiderocol (FDC) Susceptibility Testing

Iron-depleted Mueller-Hinton II broth (IDMH) was prepared according to Clinical and Laboratory Standards Institute guidelines (CLSI M100 35^th^ Ed.) using Chelex 100 resin (BioRad) to adjust iron concentrations to < 0.03 mg/L (41). For FDC broth microdilution testing, *P. aeruginosa* overnight cultures (16-20 h) were grown in IDMH. 200-fold dilution of a 0.5 McFarland standard (O.D. 600 nm 0.08 – 0.1, corresponding to ∼10^8^ CFU/mL) in IDMH was used as the starting inoculum. The assay was performed in 96-well round bottom plates (Corning) (150 µL/well) for FDC (Shionogi) concentrations 0.03 – 32 µg/mL. Minimum inhibitory concentrations (MICs) were interpreted visually after 24 h incubation at 37 °C.

FDC disk diffusion testing was performed on Mueller-Hinton II agar using 30 µg FDC disks (Hardy Diagnostics) and 0.5 McFarland standard inoculum from IDMH overnight cultures. Zones of inhibition were measured after 24 h incubation at 37 °C.

### Chrome Azurol S (CAS) Siderophore Production Assay

A modified CAS agar medium was prepared using previously established methods (42). The CAS reagent solution was prepared in 250 mL ddH_2_O (1 mM Chrome Azurol S (Honeywell Fluka), 0.1 mM FeCl_3_, 2 mM hexadecyltrimethylammonium bromide (HDTMA) (Acros Organics)) and autoclaved in a plastic container. 750 mL Mueller-Hinton II agar (BD) (concentration adjusted for 1 L final volume) was prepared and autoclaved in a separate plastic container, combined with 250 mL CAS reagent solution, and supplemented with 10 µg/mL triclosan. CAS assay was performed by spotting 10 µL of 0.5 McFarland standard inoculum in IDMH onto a 10 cm CAS agar plate. Diameter of the apo-CAS halo around the bacterial lawn was measured after 24 h and 48 h incubation at 37 °C.

### Next Generation Sequencing and Mutation Analysis

Single nucleotide polymorphisms, insertions, and deletions in PA14 gene deletion mutants or FDC-adapted strains were identified by whole genome sequencing, as previously described (43). Briefly, strains were grown at 37° C in a shaking incubator for 3-6 hours, pelleted, and genomic DNA was extracted using the DNeasy blood and tissue kit (Quiagen). Whole genome sequencing was carried out on a NextSeq2000 platform (Illumina) with 2x300 paired-end reads. Mutations were identified by aligning short-read sequences to the UCBPP-PA14 reference genome (NC_008463.1) using Bowtie2 version 2.4.5 built under the *breseq* pipeline (version 0.39.0) (44).

### Transposon Mutagenesis

Transposon mutagenesis was performed using an *E. coli* SM10 conjugal donor strain carrying the pMAR2xT7 plasmid that expresses the Mariner transposon (23). PA14 wild-type or PA14*ΔfptA* was mated with *E. coli* SM10 on TSA and transposon mutants were selected on TSA supplemented with 30 µg/mL gentamicin and 10 µg/mL triclosan. For PA14 WT, ∼50K transposon mutants were pooled into one library. For PA14*ΔfptA*, ∼50K transposon mutants were pooled into 12 sublibraries. Transposon mutant libraries were stored at -80 °C before use.

To screen transposon mutants for decreased FDC susceptibility, libraries were thawed and grown in M9 low-iron casamino acids medium (11.28 g/L Difco 5X M9 Salts (BD), 17.5 g/L Bacto Casamino Acids – low sodium chloride and iron concentrations (Gibco), 0.5 mM MgCl_2_, 0.5 mM CaCl_2_) and treated with 0.03 – 32 µg/mL FDC (1:1 M9 low-iron medium and IDMH) in a 96-well round bottom plate for 24 h at 37 °C. Mutants were isolated from the condition with the highest concentration of FDC that permitted bacterial growth. The transposon insertion sites in these mutants were identified by Sanger sequencing using arbitrary and transposon-specific primers as previously described (23).

### Quantitative Real-Time PCR (qRT-PCR)

qRT-PCR was performed to measure gene expression (mRNA) levels for CpxR-regulated genes (*muxA*, *cpxP*), genes encoding TonB-dependent siderophore receptors (*piuA*, *pirA*), and genes downstream of select transposon insertion mutants (*ponA*, *muxA*). *P. aeruginosa* was grown in M9 low-iron casamino acids medium in a 50 mL conical tube (10 mL) for 4 h at 37 °C with vigorous shaking with a starting inoculum of ∼5х10^7^ CFU/mL. RNA isolation was performed as previously described (45) using Trizol reagent (Invitrogen) according to manufacturer’s protocols. DNA contaminants were removed from the purified extract by TURBO DNase (Invitrogen) and reverse transcription was performed using a LunaScript RT Supermix (NEB). qRT-PCR was performed using PowerUp SYBR Green Master Mix (Applied Biosystems) in a CFX Opus 96 Real-Time PCR System (BioRad). Relative gene expression was calculated using a ΔΔCt method based on the values for PA14 WT and using *gyrB* as the housekeeping gene.

### Bacterial Growth and Pyoverdine Production Measurement

Bacterial growth (O.D. 600 nm) was measured using a Synergy H1 multimode microplate reader (BioTek) using a 96-well round bottom plate (with or without FDC) prepared as above. For growth curves, absorbance was measured every 30 min for 24 h with continuous incubation at 37 °C. Pyoverdine production was measured from IDMH overnight cultures by fluorescence (Ex. 405 nm; Em. 460 nm) in a 96-well black, clear flat bottom plate (Corning).

### FptA^+^/FptA^-^ Mixed Population Passaging

FptA^+^ (WT) and FptA^-^ (*ΔfptA*) mutants were generated for pyoverdine-producing (PA14*ΔpirR*) and non-producing (PA14*ΔpvdAΔpirR*) strain backgrounds. FptA^+^/FptA^-^ co-cultures were mixed at a 1:1 ratio (∼10^6^ CFU/mL each) in 2 mL IDMH and passaged in a 15 mL conical tube every 24 h for 10 days. At each passage, the co-culture was diluted 10,000-fold in IDMH and FDC disk diffusion testing was performed. Cells were harvested at the beginning (day 1) and end (day 10) of the experiment for allelic abundance analysis.

The relative abundance of each *fptA* allele in the co-culture was quantified by qPCR. Primers were designed for ∼100 bp amplification within the gene deletion region (WT-specific primers), upstream and downstream regions flanking the deletion site (*ΔfptA*-specific primers), or within the downstream region shared by both WT and *ΔfptA* strains (nonspecific primers). Genomic DNA from the co-cultures were extracted using a DNeasy UltraClean Microbial Kit (Qiagen) and adjusted to a final concentration of 100 ng/µL using a Nanodrop One spectrophotometer (ThermoFisher Scientific). qPCR reactions were performed using a PowerUp SYBR Green Master Mix in a CFX Opus 96 Real-Time PCR System. The relative abundance of the WT *fptA* and *ΔfptA* alleles in the co-culture was calculated using a ΔCt method:

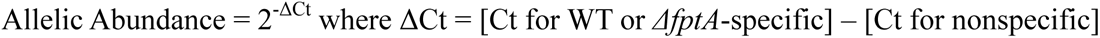

### Statistical Analysis

Student’s *t*-test and one-way ANOVA with multiple comparisons tests were performed using GraphPad Prism 10.

## Acknowledgements

This study was supported by the National Institute of Allergy and Infectious Diseases (NIH/NIAID) grants R21 AI175821 and R21 AI190338 awarded to WRM. DK was supported by a training fellowship administered by the Gulf Coast Consortia, Antimicrobial Resistance Training Program in the Texas Medical Center (AMR-TPT), that was funded by the NIH/NIAID grant T32 AI179595 and Cystic Fibrosis Foundation grant KANG25F0. CAA is supported by an NIH/NIAID grant number K24 AI121296, R01 AI148342, R01 AI134637, and P01 AI152999. We thank Dr. Stephanie Egge for constructing the *pirR* gene deletion plasmid.

## Data availability

The whole genome sequencing data is available at the National Center for Biotechnology Information (NCBI) website, Bioproject accession number PRJNA1316631.

## Transparency declarations

CAA has received royalties from UpToDate. WRM has received grant support from Merck and royalties from UpToDate. All other authors have no conflicts to disclose.

## Supplementary Figure Legends

**Figure S1. Chrome Azurol S (CAS) siderophore activity assay measures pyochelin production on Mueller-Hinton agar. (A)** Visualization of siderophore production by PA14 mutants on Mueller-Hinton agar supplemented with CAS-Fe^3+^ after 48 h. **(B)** Quantification of siderophore production using the diameter of the apo-CAS halo. * corresponds to *p* < 0.01 based on a one-way ANOVA with Dunnett’s multiple comparisons test.

**Figure S2. Mutation in *cpxS* leads to *muxA* overexpression. (A)** Alpha-fold protein structure prediction and superimposition (“Matchmaker”) for CpxS from PA14 WT and PA14*cpxS_Δ86-98_*. Structures were generated by ChimeraX. **(B)** Expression of *piuA*, *pirA*, *muxA*, and *cpxP* (by qRT-PCR) in PA14*cpxS_Δ86-98_* compared to PA14 WT. **(C)** Pyoverdine production (measured in fluorescence - Ex. 405 nm; Em. 460 nm) in iron-depleted Mueller-Hinton broth by PA14 mutants.

**Figure S3. Inactivation of *fptA* abolishes bacteriostatic effects of FDC at subinhibitory concentrations. (A-D)** Bacterial growth (O.D. 600 nm) measured every 30 min for 24 h in iron-depleted Mueller-Hinton broth with increasing concentrations of FDC (0.03 – 32 µg/mL) for PA14 WT **(A)**, PA14*ΔpirR* **(B)**, PA14*cpxS_Δ86-98_* **(C)**, and PA14*ΔfptA* **(D)**.

**Figure S4. Mixed population of *fptA^+^*and *fptA^-^* mutants is gradually resensitized to FDC in the absence of pyoverdine production. (A, B)** Images of FDC disk diffusion testing plates for co-cultures of PA14*ΔpirR* and PA14*ΔpirRΔfptA* (A) or PA14*ΔpvdAΔpirR* and PA14*ΔpvdAΔpirRΔfptA* (B) passaged in iron-depleted Mueller-Hinton broth for 10 days.

## Supplementary Tables

Table S1. Analysis of single nucleotide polymorphisms (SNPs) in PA14 mutants.

Table S2. Statistical analysis for bacterial growth curves in **Figure 3**.

Table S3. List of primers used in this study.

